# KDM6A Haploinsufficiency Is Insufficient to Disrupt Cochlear Structure and Auditory Function in Mouse Models of Kabuki Syndrome Type 2

**DOI:** 10.64898/2026.10.09.757851

**Authors:** Yuichi Shimizu, Shun Wakamori, Loryn Smith, Ghazaleh Behnammanesh, Yohei Honkura, Jun Suzuki, Jonathan Bird, Yukio Katori, Shinichi Someya

**Author notes:** Corresponding authors: Yuichi Shimizu,; Shinichi Someya.

## Abstract

**Objective:** Kabuki syndrome type 2 (KS2) is a rare X-linked disorder caused by pathogenic variants in *KDM6A* and is frequently associated with hearing impairment. Because auditory phenotypes include sensorineural, conductive, and mixed hearing loss, we examined whether *Kdm6a* deficiency directly impairs cochlear morphology or auditory function in mice.

**Methods:** We generated *Nestin-Cre; Kdm6a* conditional mutants and a germline-derived global *Kdm6a* deletion model using *Sox2-Cre*-mediated germline recombination. Auditory function and cochlear morphology were assessed by auditory brainstem responses (ABRs), distortion product otoacoustic emissions (DPOAEs), and histological analyses.

**Results:** Whole-body hemizygous or homozygous *Kdm6a* deletion resulted in embryonic lethality. Analyses therefore focused on *Nestin-Cre; Kdm6a^flox/Y^* males and females carrying a germline-derived heterozygous *Kdm6a* deletion. In *Nestin-Cre* mutants, ABR thresholds, wave I amplitudes and latencies, and DPOAE measures at 8, 16, and 32 kHz were comparable with littermate controls at 4 and 8 weeks. In heterozygous deletion females, ABR thresholds, wave I amplitudes and latencies at 4–64 kHz and DPOAE measures at 8, 16, and 32 kHz were comparable with male and female wild-type controls.

No overt cochlear pathology was observed in either model.

**Conclusion:** *Kdm6a* deficiency in the viable mouse models examined did not produce overt cochlear pathology or baseline auditory dysfunction. These findings suggest that partial *Kdm6a* loss alone is insufficient to produce a robust primary cochlear phenotype in young mice and highlight the potential contributions of non-cochlear, developmental, central, and later-onset mechanisms to hearing impairment in KS2. Clinically, these findings support careful evaluation for potentially treatable middle-ear disease.

## 1. Introduction

Hearing loss is a major global health issue affecting approximately 430 million individuals worldwide, and genetic factors account for a substantial proportion of prelingual hearing loss, particularly in developed countries [1, 2]. Recent epidemiological studies have identified hearing loss as one of the most important modifiable risk factors for dementia, accounting for approximately 8% of attributable risk and underscoring the clinical and societal impact of auditory dysfunction [3]. Despite this growing recognition, effective treatments for sensorineural hearing loss (SNHL) remain limited, with current management largely restricted to symptomatic interventions such as hearing aids and cochlear implants [4]. Because SNHL is generally irreversible, a better understanding of its underlying molecular and genetic mechanisms is essential for the development of disease-modifying therapies.

Kabuki syndrome (KS) is a rare multisystem genetic disorder characterized by distinctive craniofacial features and a broad spectrum of congenital and neurodevelopmental manifestations, including growth retardation, skeletal anomalies, and immune dysfunction [5]. Hearing impairment is commonly reported in individuals with KS, affecting approximately 30% of patients [5]. However, the auditory phenotype of KS remains incompletely characterized, and the underlying pathophysiologic mechanisms are not well understood. Published studies describe considerable heterogeneity in both the severity and type of hearing impairment, including sensorineural, conductive, and mixed hearing loss [6–9], suggesting that multiple mechanisms may contribute to auditory dysfunction in KS.

Most cases of KS are caused by pathogenic variants affecting one of two epigenetic regulators: *KMT2D*, which causes Kabuki syndrome type 1 (KS1), and *KDM6A*, which causes Kabuki syndrome type 2 (KS2) [10]. Pathogenic variants in *KMT2D* account for approximately 56–76% of cases, whereas pathogenic variants in *KDM6A* account for approximately 5–8% [10]. *KDM6A* is X-linked, and KS2 is typically associated with heterozygous loss-of-function variants in females or hemizygous variants in males, with variable phenotypic expressivity [11]. The absence of reported individuals with complete biallelic loss of KDM6A [11], together with findings from animal studies [12], suggests that complete loss of KDM6A may result in severe developmental failure or embryonic lethality.

KDM6A, also known as lysine demethylase 6A, catalyzes the removal of the repressive histone mark H3K27me3, thereby regulating chromatin accessibility and gene transcription [13]. KDM6A plays essential roles in embryonic development, cell fate determination, and tissue differentiation and has also been implicated in tumor suppression [14]. Single-cell RNA sequencing studies have demonstrated *Kdm6a* expression across multiple inner ear cell types [15], raising the possibility that KDM6A may play a direct role in cochlear development and auditory function. In addition, *Kdm6a* consistently escapes X-chromosome inactivation [16], resulting in higher expression levels in females than in males. This sex-biased expression, together with the X-linked inheritance of KS2, highlights the importance of defining how reduced *Kdm6a* dosage affects cochlear structure and function.

Although hearing impairment is frequently reported in individuals with KS2, direct experimental evidence linking KDM6A haploinsufficiency to primary cochlear dysfunction remains limited. It therefore remains unclear whether reduced KDM6A dosage is sufficient to produce a primary cochlear phenotype. Resolving this question is important for interpreting the heterogeneous auditory phenotypes observed in KS2 and for distinguishing potential cochlear contributions from conductive, developmental, central, or other mechanisms of hearing impairment. To address this knowledge gap, we generated and characterized complementary *Kdm6a* deletion mouse models and performed genetic, physiological, and histological analyses to determine whether partial *Kdm6a* loss is sufficient to disrupt cochlear structure and auditory function.

## 2. Materials and Methods

### 2.1. Animals

All mouse strains used in this study were obtained from The Jackson Laboratory (Bar Harbor, ME, USA). These included C57BL/6J mice (strain #: 000664), *Kdm6a* floxed mice (strain #: 024177), *Sox2-Cre* transgenic mice (strain #: 008454), and *Nestin-Cre* mice (strain #: 003771). Nestin promoter-driven Cre recombinase is expressed in the central and peripheral nervous systems, including neuronal and glial precursor cells beginning during embryonic development [17]. We used this line to generate neural-lineage *Kdm6a* conditional mutants; however, the precise pattern of Cre-mediated recombination within the peripheral and central auditory pathways was not independently mapped in this study. In contrast, because *Sox2-Cre* transgene expression is active in the female germ line, *Sox2-Cre* mice were used to generate germline-derived, whole-body *Kdm6a* deletion models through germline recombination.

Because *Kdm6a* is located on the X chromosome, conditional mutant genotypes were generated in a sex-dependent manner by crossing *Kdm6a* floxed mice with Cre-driver lines. For the *Nestin-Cre* line, *Nestin-Cre* homozygous male mice were first crossed with *Kdm6a^flox/flox^* female mice to generate *Nestin-Cre/+*; *Kdm6a^flox/Y^* male conditional mutants. These male mice were subsequently crossed with *Kdm6a^flox/flox^* females to generate *Nestin-Cre/+*; *Kdm6a^flox/Y^* male mice as well as *Nestin-Cre/+; Kdm6a^flox/flox^ and Nestin-Cre/+; Kdm6a^flox/+^* female conditional mutants. For the *Nestin-Cre* line, littermate controls were defined as mice carrying the corresponding floxed allele but lacking the *Nestin-Cre* transgene.

For the *Sox2-Cre* line, *Sox2-Cre* mice were used for germline recombination because Cre activity in this line is present in the female germ line, enabling whole-body deletion of floxed alleles in the offspring, as previously described [18]. *Sox2-Cre* heterozygous mice were crossed with male *Kdm6a^flox/Y^* mice to generate females carrying the germline-derived *Kdm6a^del/WT^* allele. These females were subsequently crossed with *Kdm6a^flox/Y^* male mice to assess the expected genotypes, including *Kdm6a^del/Y^*, *Kdm6a^del/del^*, and *Kdm6a^del/WT^* mice. Among the heterozygous females used for phenotypic analyses, those used in Fig. 4 carried the *Kdm6a^del/WT^*allele without the *Sox2-Cre* transgene, whereas those used in Supplementary Fig. 2 carried the *Kdm6a^del/WT^* allele together with the *Sox2-Cre* transgene (*Tg/+*). For the germline-derived deletion model, wild-type male and female mice were used as controls.

All animal experiments were conducted under protocols approved by the Institutional Animal Care and Use Committee of the institution where the experiments were performed. Mice were maintained in a temperature-controlled environment under a 12 h light/12 h dark cycle (08:00–20:00) at 23 °C with free access to food and water.

### 2.2. Genotyping

Genomic DNA was isolated from mouse tail biopsies and used for PCR-based genotyping.

For *Kdm6a* floxed mice, primer sequences were as follows: forward 5′-GCTACTGGGGTGTTTTGAATG-3′ and reverse 5′-TTTCATAGAACAGTTTCAGGATACC-3′. Cycling conditions were 95 °C for 2 min; 35 cycles of 95 °C for 30 s, 65 °C for 1 min, and 72 °C for 1 min; and 72 °C for 5 min. Expected PCR product sizes were 249 bp for the wild-type allele and 430 bp for the floxed allele.

For *Nestin-Cre* genotyping, the assay was performed according to the Jackson Laboratory protocol. Primers used were mChr12 wild type forward 5′-TTGCTAAAGCGCTACATAGGA-3′, mChr12 common reverse 5′-GCCTTATTGTGGAAGGACTG-3′, and hGH1 transgene forward 5′-CCTTCCTGAAGCAGTAGAGCA-3′. The wild type allele yielded a 246-bp band, whereas the *Nestin-Cre* transgene yielded a 150-bp band.

For *Sox2-Cre* genotyping, primers used were mChr13 wild type forward 5′-CTTGTGTAGAGTGATGGCTTGA-3′, mChr13 common reverse 5′-TAGTGCCCCATTTTTGAAGG-3′, and transgene forward 5′-CCAGTGCAGTGAAGCAAATC-3′. Cycling conditions were 95 °C for 2 min; 35 cycles of 95 °C for 30 s, 65 °C for 30 s, and 72 °C for 30 s; and 72 °C for 5 min. Expected band sizes were 207 bp and 165 bp.

For confirmation of the deleted *Kdm6a* allele, primers were used as previously described [12]: forward 5′-AACAAAAACCCAGGCTTTATTCAC-3′ and reverse 5′-AGTTTCAGGATACCTTTACTATAAG-3′. Cycling conditions were 95 °C for 2 min; 35 cycles of 95 °C for 30 s, 50 °C for 30 s, and 72 °C for 1 min 10 s; and 72 °C for 5 min. Expected PCR product sizes were 741 bp for the wild type allele and 361 bp for the deleted allele. PCR products were separated on 2% agarose gels.

### 2.3. Quantitative RT-PCR

Total RNA was extracted from whole inner ear tissues including the bony labyrinth, organ of Corti, lateral wall/stria vascularis, spiral ganglion, and vestibule using the RNeasy Lipid Tissue Mini Kit (QIAGEN, 74804). cDNA was synthesized from total RNA using the SuperScript III First-Strand Synthesis System (Thermo Fisher Scientific, 18080051) according to the manufacturer’s instructions. Quantitative PCR was performed using iTaq Universal SYBR Green Supermix (Bio-Rad, 1725120) on a CFX96 Touch Real-Time PCR Detection System (Bio-Rad). Relative expression levels of *Kdm6a* were normalized to *Actb*. The following primers were used: *Kdm6a* forward 5′-AGGAAGCAGAACACCAGGTC-3′, *Kdm6a* reverse 5′-CAGTAGCCTTCAGGAACAACAAAC-3′; *Actb*forward 5′-AGCCATGTACGTAGCCATCCA-3′, *Actb* reverse 5′-TCTCCGGAGTCCATCACAATG-3′. The *Kdm6a* primer pair was designed to amplify a region containing exon 24, such that transcripts lacking exon 24 would not be amplified. Sample sizes for each experiment are indicated in the corresponding figure legends.

### 2.4. RNA in situ hybridization

RNA in situ hybridization was performed using the BaseScope RED assay (Advanced Cell Diagnostics) according to the manufacturer’s instructions with minor modifications [19]. Cochleae were perfusion-fixed through the round and oval windows using 4% paraformaldehyde (PFA) in PBS and immersed in 4% PFA for 3 h at room temperature, and dissected in 20% RNAlater (Invitrogen). Isolated organ of Corti samples were pretreated by removing RNAlater and replaced with 200 µL of 50% ethanol and incubated for 5 min at room temperature. Samples were taken through a graded ethanol series (70%, 95%, 100%) for 5 min each. Ethanol was then aspirated and pre-treatment 1 (hydrogen peroxide, ACD) and incubated for 30 min at room temperature. Samples were then washed twice in MilliQ water for 3 min each. After the final wash, samples were incubated with a 1:100 dilution of a low-pH antigen unmasking solution (H-3300, Vector Laboratories) in MilliQ water, for 5 min at 65 °C. Samples were washed twice in MilliQ water. Following the last wash, samples were incubated with Protease Plus (ACD) for 30 min at room temperature. Samples were washed with MilliQ water and stored at 4 °C overnight. The following day, probes were added to each sample and incubated at 40 °C in a humidified oven for 2 hours. *Kdm6a* targeting probes were designed by ACD; positive control (BA-Mm-Ppib-1ZZ), negative control (BA-DapB-1ZZ). After hybridization, tissues were washed 3 times with 1x wash buffer from the RNAscope wash buffer reagents (#320058, ACD). Samples were then sequentially incubated with amplification reagents 1–8 from the BaseScope v2-RED (#323910, ACD Bio) at 40 °C for 30 min. Samples were washed three times in 1x wash buffer between amplification steps. A RED working solution was prepared just before use by mixing BaseScope Fast RED-B and BaseScope Fast RED-A at a 1:60 ratio. Samples were incubated in RED solution for 10 min at room temperature, followed by 3 x 5 min washes in MilliQ water. Samples were then incubated with a mouse monoclonal anti-MYO7A antibody (1:100, Santa Cruz Biotechnology, sc-74516) diluted in 0.1% Triton X-100 in PBS overnight at 4 °C. Following washes in PBS, samples were incubated with Alexa Fluor 488 Goat anti-mouse IgG2a (1:200, A-21131, Thermo Fisher Scientific) in PBS with 0.1% Triton X-100 for 2 hours at room temperature. After 3 times for 3 min washes in PBS, samples were incubated with 1.5 µg/mL 4’,6-diamidino-2-phenylindole (DAPI) diluted in PBS for 15 min at room temperature. After 3 washes with PBS, tissues were mounted in ProLong Glass (P36980, Invitrogen) and imaged using a confocal spinning disk microscope with a 100x oil immersion objective lens. Z stacks were captured at 0.3 μm spacing.

### 2.5. Cochlear immunofluorescence

Mice were euthanized by cervical dislocation and temporal bones were harvested. Cochleae were perfusion-fixed through the round and oval window using 4% paraformaldehyde (PFA) in PBS and immersed in 4% PFA for 2 h at room temperature, decalcified in 0.12 M EDTA in PBS for 2 days, and microdissected. Samples were blocked with 20% normal goat serum and 0.3% Triton X-100 in PBS for 1 h at room temperature, followed by overnight incubation at 4 °C with primary antibodies: mouse anti-MYO7A (1:200, Santa Cruz Biotechnology, sc-74516), rabbit anti-KDM6A (1:500, Novus, NBP1-80628), rabbit anti-TUBB3 (beta III Tubulin) (1:1000, Abcam, 18207). For the experiment shown in Fig. 4B, the blocking step was omitted, and samples were incubated directly with primary antibodies after washing. After washing, samples were incubated with appropriate fluorescent secondary antibodies for 1 h, counterstained with DAPI, and mounted with VECTASHIELD (Vector Laboratories). Images were acquired using a Nikon Ti2 inverted microscope equipped with a Yokogawa CSU-X1 spinning disk confocal scanner and analyzed using NIS-Elements and ImageJ.

### 2.6. Cochlear immunohistochemistry

Mouse inner ears were dissected and perfusion-fixed through the round and oval windows using 10% formaldehyde. Samples were then immersed in 10% formaldehyde in a 30-mm dish or 1.5-mL tube and fixed overnight. The following day, samples were transferred to 10% EDTA in a 1.5-mL tube and decalcified for 2–3 days. After decalcification, the specimens were transferred to 70% ethanol, and dehydrated for 1 day before paraffin embedding. Paraffin-embedded sections were deparaffinized and washed in distilled water. Antigen retrieval was performed using antigen retrieval buffer (pH 6), followed by cooling and rinsing in distilled water. Sections were then washed in phosphate-buffered saline (PBS) and incubated with a rabbit anti-KDM6A primary antibody (1:500, Novus, NBP1-80628) overnight at 4 °C. After washing in PBS, endogenous peroxidase activity was quenched for 20 min at room temperature using 0.3% hydrogen peroxide in methanol. Sections were then washed in PBS and incubated with Histofine Simple Stain Mouse MAX-PO (Rabbit, Nichirei) as the secondary antibody for 40 min at room temperature. After additional PBS washes, immunoreactivity was visualized using 3,3′-diaminobenzidine (DAB) solution prepared in 0.05 M Tris buffer, and the reaction was monitored microscopically for approximately 5 min. Sections were subsequently washed in PBS, rinsed in running water and distilled water, and counterstained with hematoxylin for 50 s. Finally, sections were dehydrated, cleared, and coverslipped.

### 2.7. Body weight and body length

Body weight was measured weekly, and body length was measured monthly in experimental mice and littermate controls. Measurements obtained at 4 and 8 weeks of age were collected from the same cohort of mice used for ABR, DPOAE, and rotarod testing.

### 2.8. ABR

Auditory brainstem response (ABR) thresholds were measured using a TDT system (Tucker-Davis Technologies) as previously described [20, 21]. Mice were anesthetized with ketamine (100 mg/kg) and xylazine (10 mg/kg) by intraperitoneal injection. Subdermal needle electrodes were placed at the vertex (active), ipsilateral ear (reference), and contralateral ear (ground). ABR waveforms were collected in response to tone bursts at 8, 16, and 32 kHz in *Nestin-Cre*; *Kdm6a flox* mice, and at 4, 8, 16, 32, 48 and 64 kHz in *Sox2-Cre*; *Kdm6a flox* mice. At each frequency, the sound level was reduced in 5–10 dB steps from 90 to 10 dB SPL. The ABR threshold was defined as the lowest stimulus level at which the ABR wave I was reproducibly identified. Latencies and amplitudes for ABR wave I were measured at 80 dB SPL. The ABR wave I latency was determined by measuring the amount of time elapsed from the onset of the stimulus to the highest value (peak) of ABR wave I. The ABR wave I amplitude was determined by measuring the voltage difference between the highest value (peak) and the lowest value (trough) of ABR wave I. ABR testing was performed longitudinally at 4 and 8 weeks of age in the same cohort of mice used for DPOAE, rotarod, and body measurements.

### 2.9. DPOAE

Distortion product otoacoustic emissions (DPOAE) were measured using the TDT RZ6 system with BioSigRZ software (Tucker-Davis Technologies) as previously described [22]. Mice were anesthetized with ketamine (100 mg/kg) and xylazine (10 mg/kg). Two primary tones (f1 and f2) with an f2/f1 ratio of 1.2 were presented at frequencies centered at 8, 16, and 32 kHz. Stimulus levels were reduced from 80 to 20 dB SPL. The noise floor was defined as the average of the distortion products at ten neighboring frequencies (five above and five below the 2f1-f2) [23]. The DPOAE threshold was defined as the lowest stimulus level at which the 2f1-f2 response exceeded the noise floor by at least 6 dB [23]. DPOAE testing was performed longitudinally at 4 and 8 weeks of age in the same cohort of mice used for ABR, rotarod, and body measurements.

### 2.10. Rotarod performance test

Rotarod performance was assessed using a rotarod apparatus (Rota Rod Rotamex 5; Columbus Instruments) as previously described [20]. Mice were trained using accelerating trials (4–40 rpm) during the first five trials. Performance was assessed as the average latency to fall across trials 6–8. The same cohort of mice was tested longitudinally at 4 and 8 weeks of age.

### 2.11. Statistical analysis

Data were analyzed using GraphPad Prism version 10.0.2. For longitudinal measures obtained from the same animals at 4 and 8 weeks of age (ABR, DPOAE, rotarod, and body measurements), genotype × time effects were evaluated using two-way repeated-measures ANOVA or a mixed-effects model, with Bonferroni correction for multiple comparisons, as appropriate. For within-animal comparisons between 4 and 8 weeks, paired two-tailed Student’s *t* tests were used. For unpaired comparisons (e.g., RT-qPCR), unpaired two-tailed Student’s *t* tests were used. A *P* value < 0.05 was considered statistically significant.

## 3. Results

### 3.1. *Kdm6a* is Broadly Expressed in the Inner Ear

We first examined whether KDM6A is expressed in inner ear tissues of wild-type mice using C57BL/6J mice. RT–qPCR analysis showed that *Kdm6a* mRNA was expressed in inner ear tissues of both male and female mice (Fig. 1A). *Kdm6a* mRNA levels were approximately twofold higher in female inner ears than in male inner ears, consistent with *Kdm6a* being an X-linked gene. These findings further demonstrate that *Kdm6a* escapes X-chromosome inactivation (XCI) in the mouse inner ear, resulting in higher expression levels in females.

**Figure 1.**
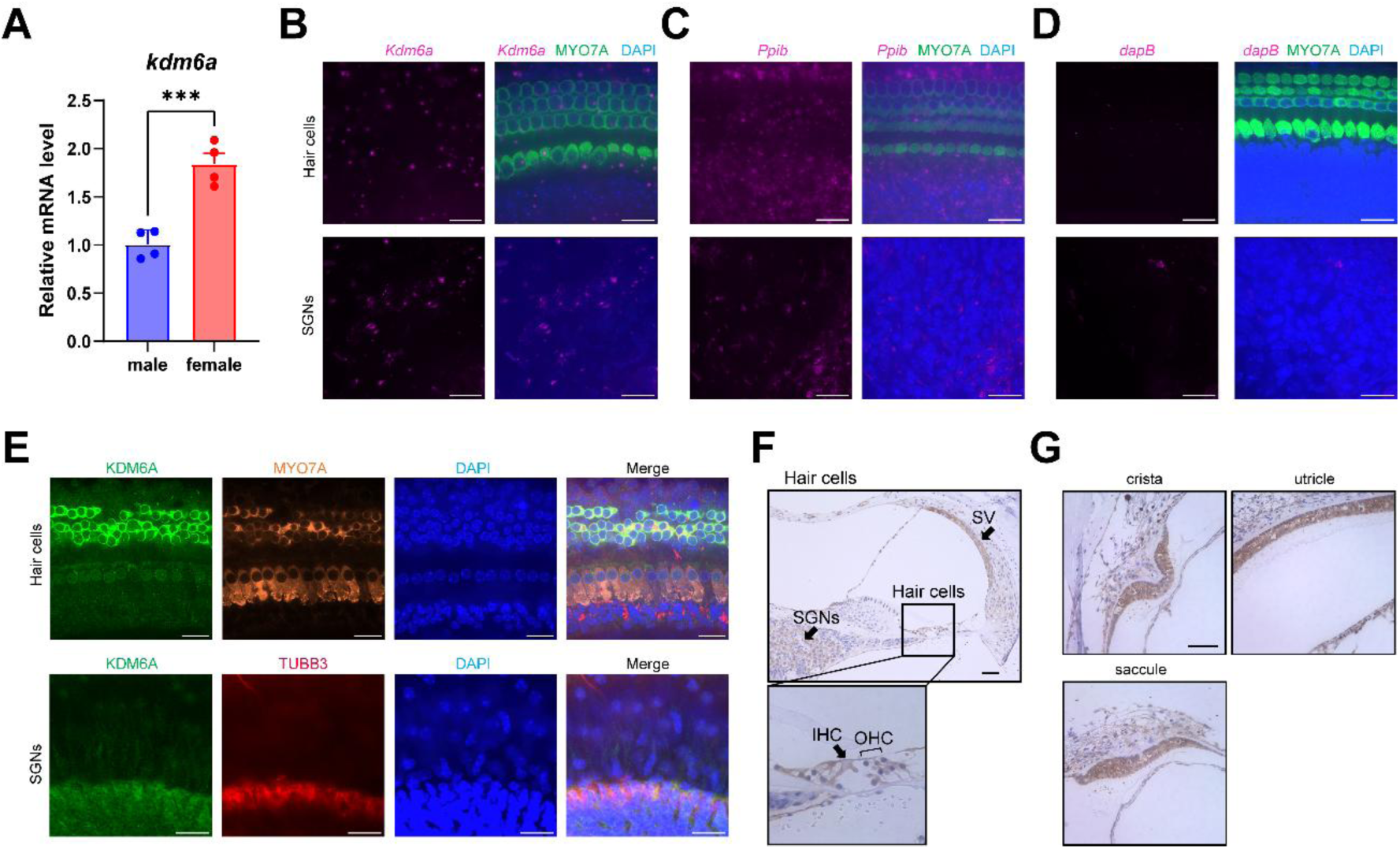
*Kdm6a* Is Broadly Expressed in the Inner Ear and Escapes X-Chromosome Inactivation. **(A)** *Kdm6a* expression in whole inner ear tissue from male and female C57BL/6J mice aged 5–16 weeks, as measured by RT– qPCR. Each dot represents an individual animal (n = 4 per sex). Bars indicate mean ± SEM. **(B)** BaseScope in situ hybridization showing *Kdm6a* signals (magenta) in cochlear whole mounts co-stained for MYO7A (green) and DAPI (blue). *Kdm6a* signals were detected in inner hair cells (IHCs), outer hair cells (OHCs), supporting cells, and auditory nerve fibers (ANs) (left panel), as well as in spiral ganglion neurons (SGNs) (right panel). **(C)** Positive control with the mouse *Ppib* probe. *Ppib*, peptidylprolyl isomerase B. **(D)** Negative control using the bacterial *dapB* probe. *dapB*, 4-hydroxy-tetrahydrodipicolinate reductase. The *Kdm6a* and *dapB* images were obtained from male mice, whereas the *Ppib* image was obtained from a female mouse. All mice shown in panels B–D were examined at postnatal day 5 (P5). Scale bars, 20 µm. **(E)** Immunofluorescence images of cochlear whole mounts from a 5-week-old male mouse stained for KDM6A (green), MYO7A (orange), TUBB3 (red) and DAPI (blue). KDM6A immunoreactivity was detected in OHCs, IHCs, supporting cells, ANs, and SGNs. Scale bars, 20 µm. **(F)** Paraffin section of the cochlea from a 12-week-old male mouse stained for KDM6A by immunohistochemistry. KDM6A immunoreactivity (brown) was detected in hair cells, supporting cells, SGNs, and the stria vascularis (SV). Scale bar, 50 µm. **(G)** Paraffin sections of the vestibular organs from the same mouse, stained for KDM6A by immunohistochemistry. KDM6A immunoreactivity (brown) was detected in the crista, utricle, and saccule (arrows). Scale bars, 50 µm. ***P < 0.001.

We next examined the cellular distribution of *Kdm6a*/KDM6A in the inner ear. In cochlear whole mounts, BaseScope RNA in situ hybridization demonstrated *Kdm6a* mRNA expression in inner hair cells, outer hair cells, supporting cells, auditory nerve fibers, and spiral ganglion neurons (Fig. 1B–D). Consistent with these findings, immunofluorescence staining showed KDM6A protein expression in outer hair cells, with moderate expression detected in inner hair cells, auditory nerve fibers, and spiral ganglion neurons (Fig. 1E). Using paraffin-embedded sections, we further confirmed KDM6A expression in the cochlea and vestibular organs. KDM6A-positive signals were detected in hair cells, supporting cells, spiral ganglion neurons, and the stria vascularis, with relatively strong staining in the stria vascularis (Fig. 1F). KDM6A expression was also observed in vestibular sensory organs, including the crista, utricle, and saccule (Fig. 1G). These findings indicate that *Kdm6a*/KDM6A is broadly expressed throughout the inner ear, with cell-type-specific differences in staining intensity.

### 3.2. Generation of *Kdm6a* Conditional Knockout Mouse Lines

To examine whether *Kdm6a* deficiency directly affects auditory function, we generated two mouse lines: *Nestin-Cre; Kdm6a flox* mice and *Sox2-Cre; Kdm6a flox* mice. Because *Kdm6a* is located on the X chromosome, the genotypes of the conditional mutants were sex dependent.

First, we generated *Nestin-Cre; Kdm6a flox* conditional mutant mice, in which *Kdm6a* was expected to be deleted in the central and peripheral nervous systems, including neuronal and glial precursor cells [17]. In the *Kdm6a* floxed allele, loxP sites flank exon 24, and Cre-mediated recombination excises exon 24 to generate a deleted allele (Fig. 2A). Genotyping PCR confirmed the presence of the *Nestin-Cre* transgene as a 150-bp band and the wild-type allele as a 246-bp band (Fig. 2B). Genotyping PCR for the *Kdm6a* floxed allele identified the floxed and wild-type alleles as 430-bp and 249-bp bands, respectively (Fig. 2C). Using this strategy, we obtained both male and female conditional mutant mice.

**Figure 2.**
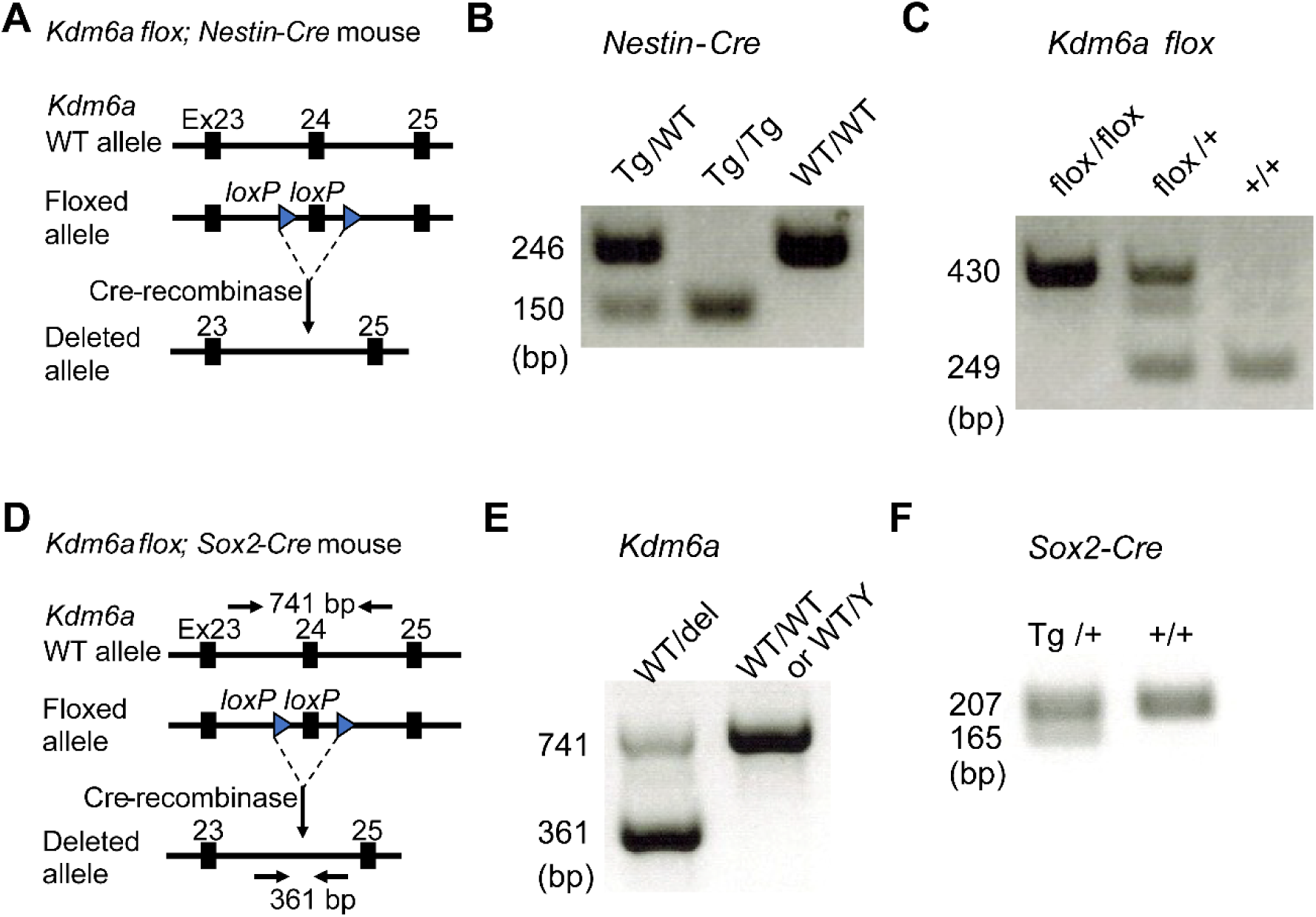
Generation of *Kdm6a* Deletion Mouse Models. **(A)** Schematic diagram of the *Kdm6a* floxed allele used to generate the *Nestin-Cre* conditional mutant model. Exon 24 is flanked by loxP sites and is deleted after Cre-mediated recombination. **(B)** Genotyping PCR for the *Nestin-Cre* transgene using tail DNA. The wild-type allele yields a 246-bp band, whereas the *Nestin-Cre* transgene yields a 150-bp band. **(C)** Genotyping PCR for the *Kdm6a* floxed allele. The floxed allele yields a 430-bp band, and the wild-type allele yields a 249-bp band. **(D)** Schematic diagram of the *Kdm6a* floxed allele used to generate the *Sox2-Cre* germline-derived deletion model, showing the expected PCR products after Cre-mediated recombination. The wild-type allele yields a 741-bp band, whereas the deleted allele yields a 361-bp band. **(E)** PCR-based confirmation of the deleted *Kdm6a* allele in genomic DNA following *Sox2-Cre*– mediated germline recombination.. **(F)** Genotyping PCR for the *Sox2-Cre* transgene. The transgene-positive allele yields a 165-bp band, whereas the wild-type allele yields a 207-bp band.

Next, to generate whole-body *Kdm6a* conditional knockout mice, we generated *Sox2-Cre; Kdm6a flox* mice. In the *Sox2-Cre* line, Cre recombinase is expressed in the germ line and is therefore expected to mediate deletion of exon 24 in all cells. Genotyping PCR for the deleted allele was designed to yield a 741-bp band for the wild-type allele and a 361-bp band for the deleted allele (Fig. 2D, E). In addition, genotyping PCR for the *Sox2-Cre* transgene identified a 165-bp band for the transgene-positive allele and a 207-bp band for the wild-type allele (Fig. 2F). Although we attempted to generate *Kdm6a^del/Y^* and *Kdm6a^del/del^*mice using this strategy, no hemizygous male or homozygous female mutants were obtained.

In both the *Nestin-Cre* and *Sox2-Cre* lines, the relevant alleles and Cre transgenes were confirmed by genotyping PCR using tail samples.

### 3.3. *Nestin-Cre; Kdm6a* Conditional Mutants Show Reduced Brain *Kdm6a* Expression but Preserved Cochlear KDM6A Expression and Auditory Function

To examine the effects of conditional knockout of *Kdm6a* in the central nervous system and the peripheral nervous system, we first generated *Nestin-Cre*; *Kdm6a* floxed conditional mutant mice, in which *Kdm6a* was expected to be deleted in neuronal and glial lineages. Because *Nestin-Cre; Kdm6a^flox/flox^* female mice were not viable, subsequent analyses focused on *Nestin-Cre; Kdm6a^flox/Y^* male mice. RT–qPCR analysis demonstrated significantly lower *Kdm6a* expression in brain tissues from *Nestin-Cre* conditional mutants compared with littermate controls. In contrast, *Kdm6a* expression in the whole inner ear was significantly higher in *Nestin-Cre* conditional mutants than in littermate controls (Fig. 3A). Consistent with the absence of reduced inner ear *Kdm6a* expression, immunostaining revealed persistent KDM6A protein expression in the cochleae of *Nestin-Cre* conditional mutants (Fig. 3B). Together, these findings demonstrate reduced *Kdm6a* expression in bulk brain tissue but no corresponding reduction in inner ear *Kdm6a* mRNA expression or loss of cochlear KDM6A immunoreactivity.

**Figure 3.**
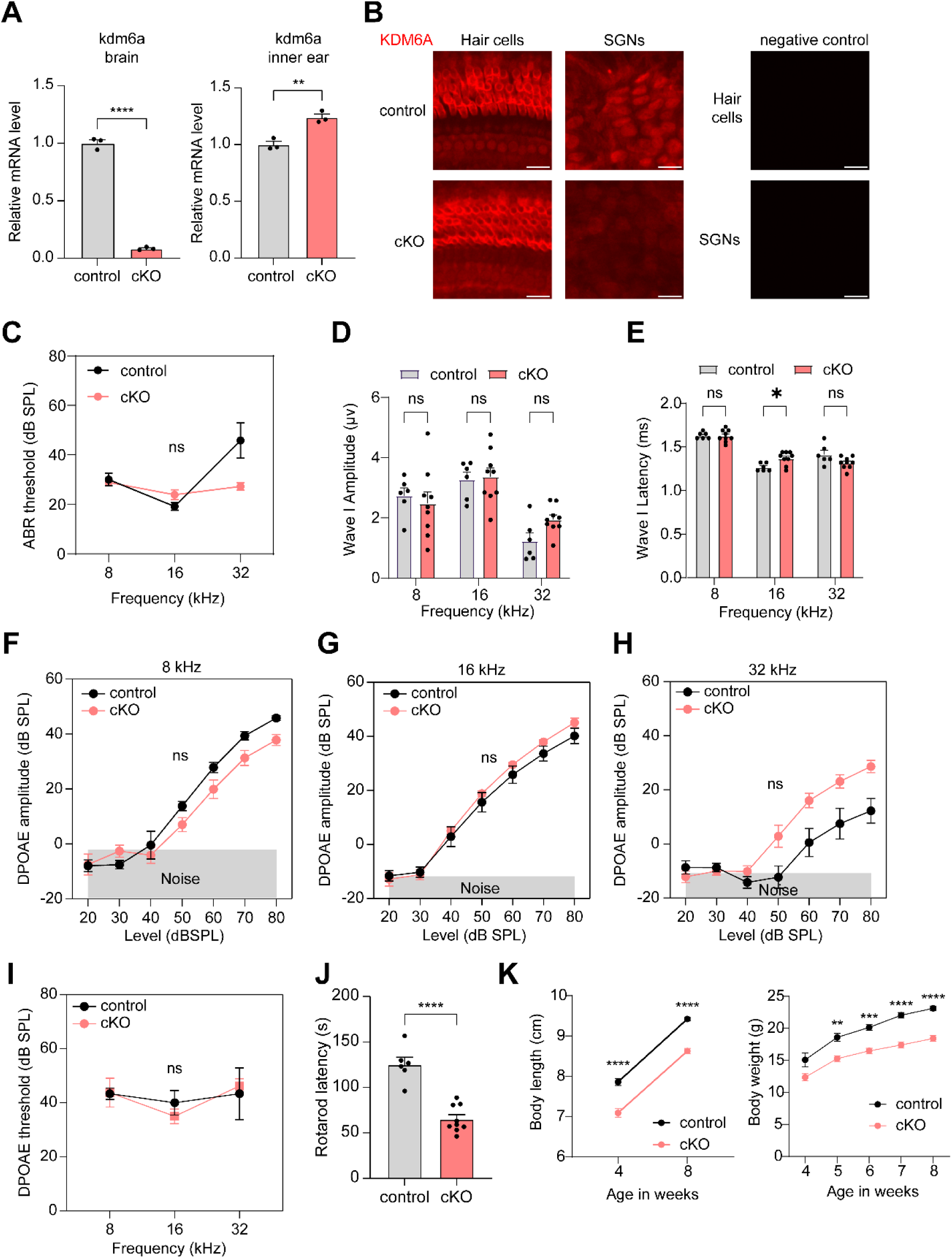
*Nestin-Cre*; *Kdm6a* Conditional Mutants Show Reduced Brain *Kdm6a* Expression but Preserved Cochlear Expression and Auditory Function. (A) *Kdm6a* expression in brain and whole inner ear tissue from 3-week-old male control and *Nestin-Cre*; *Kdm6a* conditional mutant (cKO) mice, as measured by RT–qPCR. n = 3 per group; bars indicate mean ± SEM. (B) Immunofluorescence images of cochlear whole mounts from control and *Nestin-Cre*; *Kdm6a* conditional mutant (cKO) mice at 10 weeks of age, stained for KDM6A. Images from negative controls processed without the primary antibody are also shown. Scale bars, 20 μm. **(C–E)** Auditory function assessed by ABR thresholds, wave I amplitudes, and wave I latencies in male control and cKO mice at 8 weeks of age. n = 6 in control and n = 9 in cKO; bars indicate mean ± SEM. **(F–H)** DPOAE amplitudes at 8, 16, and 32 kHz in male control and cKO mice at 8 weeks of age. n = 6 in control and n = 8 in cKO; bars indicate mean ± SEM. **(I)** DPOAE thresholds in male control and cKO mice at 8 weeks of age. **(J)** Rotarod performance in male control and cKO mice at 8 weeks of age. n = 6 in control and n = 9 in cKO; bars indicate mean ± SEM. **(K)** Longitudinal body length and body weight measurements from 4 to 8 weeks of age in the same animals; n = 6–7 in control and n = 9–11 in cKO; bars indicate mean ± SEM. *p < 0.05, **p < 0.01, ***p < 0.001, and ****p < 0.0001; ns, not significant.

Although a small difference in ABR wave I latency was observed at 16 kHz (Fig. 3C–E), *Nestin-Cre* conditional mutants otherwise exhibited preserved auditory function. At 8 weeks of age, ABR thresholds, wave I amplitudes, and wave I latencies were comparable between *Nestin-Cre* conditional mutants and littermate controls, consistent with the absence of reduced *Kdm6a*/KDM6A expression in the cochlea. Similarly, DPOAE responses were preserved (Fig. 3F–I). A similar auditory phenotype was observed at 4 weeks of age (Supplementary Fig. 1A–G). In contrast to the preserved auditory phenotype, *Nestin*-*Cre* conditional mutants exhibited impaired rotarod performance, with a shorter latency to fall than littermate controls at both 4 and 8 weeks of age (Fig. 3J, Supplementary Fig. 1H). Conditional mutants also displayed reduced body weight and shorter body length compared with age-matched controls (Fig. 3K). These systemic and motor phenotypes occurred in parallel with reduced *Kdm6a* expression in brain tissue; however, their underlying neuroanatomical basis was not determined.

### 3.4. Whole-Body Heterozygous *Kdm6a* Deletion Is Insufficient to Impair Cochlear Morphology or Auditory Function

Because *Nestin*-*Cre*–mediated hemizygous *Kdm6a* deletion did not eliminate *Kdm6a* expression in inner ear tissues, we generated mice with whole-body *Kdm6a* deletion using *Sox2*-*Cre*–mediated germline recombination. Hemizygous male and homozygous female mice with global *Kdm6a* deletion (*Kdm6a^del/Y^* and *Kdm6a^del/del^,* respectively) were not recovered among viable offspring and therefore could not be evaluated. Subsequent analyses focused on heterozygous female mice together with wild-type male and female littermate controls.

We first examined KDM6A protein expression in cochlear whole mounts from wild-type male, heterozygous female, and wild-type female mice. In outer hair cells, KDM6A immunoreactivity was most prominent along the cell periphery, with weaker cytoplasmic signals and minimal nuclear staining (Fig. 4A–H). In contrast, inner hair cells exhibited more prominent nuclear and cytoplasmic KDM6A immunoreactivity. Weaker KDM6A signals were also detected in supporting cells and auditory nerve fibers, whereas spiral ganglion neurons displayed predominantly nuclear and perinuclear staining. Qualitatively, KDM6A immunoreactivity appeared to increase in the following order: wild-type male < heterozygous female < wild-type female (Fig. 4A–C, E–G), consistent with reduced KDM6A protein expression in heterozygous females compared with wild-type females. No overt cochlear pathology was observed in heterozygous female mice.

**Figure 4.**
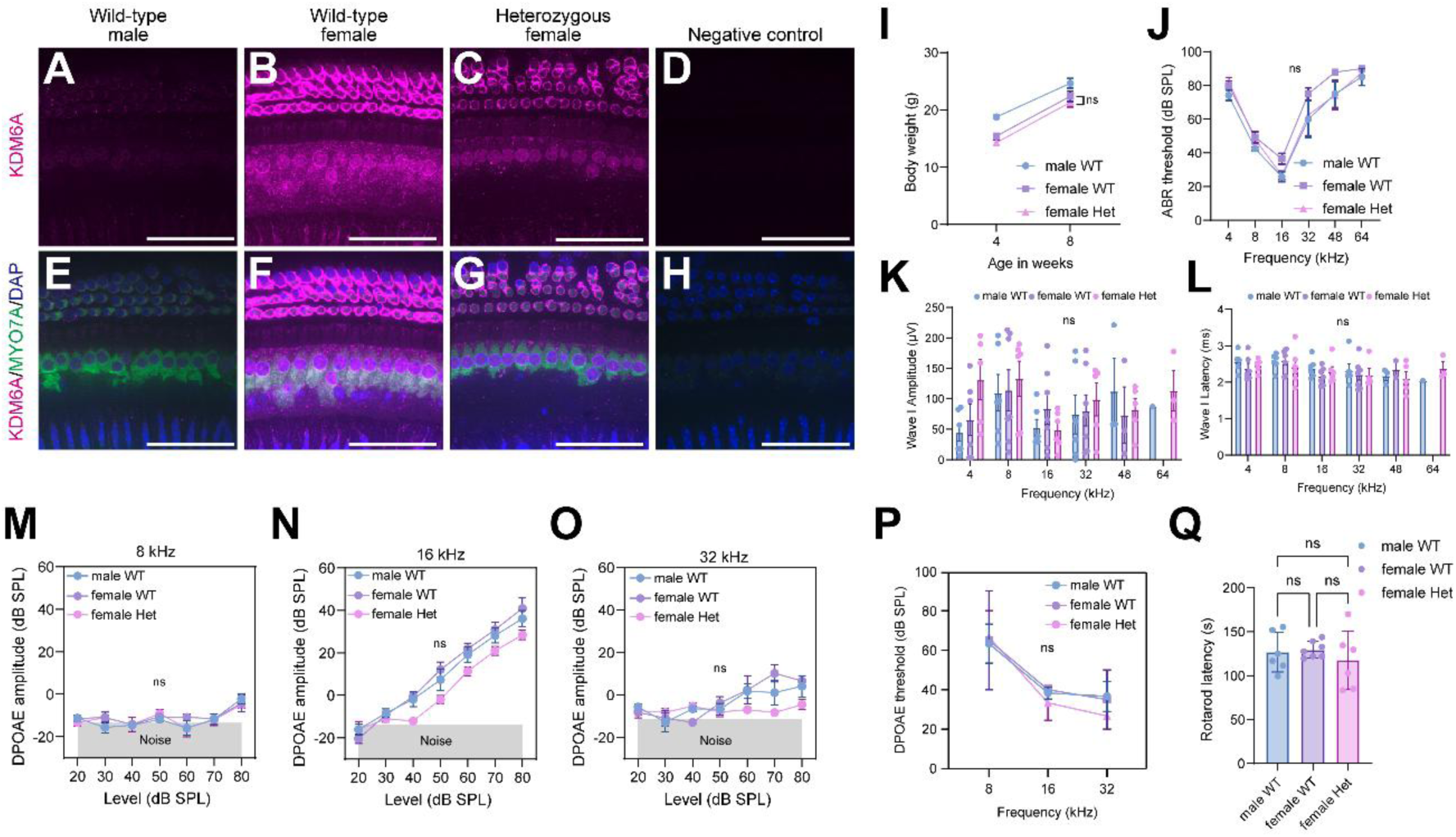
Germline-Derived Heterozygous *Kdm6a* Deletion Is Insufficient to Impair Cochlear Morphology or Auditory Function. **(A–H)** Cochlear whole-mount preparations from 9 to 11-week-old mice were immunostained for KDM6A (magenta), MYO7A (green), and DAPI (blue). Images were obtained from the apical turn using a 100× objective. **(A–D)** KDM6A-channel images from wild-type male, wild-type female, heterozygous female, and negative-control samples, respectively. **(E–H)** Corresponding merged images. Scale bars, 20 μm. **(I)** Body weight at 4 and 8 weeks of age. No significant differences were observed between wild-type and heterozygous females. **(J–L)** Auditory function assessed by ABR thresholds (**J**), wave I amplitudes (**K**), and wave I latencies (**L**) at 8 weeks of age. **(M–O)** DPOAE amplitudes at 8, 16, and 32 kHz at 8 weeks of age. **(P)** DPOAE thresholds at 8 weeks of age. **(Q)** Rotarod performance at 8 weeks of age. For panels **I**–**Q**, n = 6 for wild-type males, n = 6 for heterozygous females, and n = 7 for wild-type females; bars indicate mean ± SEM. ns, not significant.

Heterozygous female mice appeared grossly normal, and body weight at 8 weeks of age did not differ between heterozygous and wild-type female mice (Fig. 4I). Auditory function was also preserved in heterozygous females. At 8 weeks of age, ABR thresholds, wave I amplitudes and latencies across 4–64 kHz were comparable among wild-type male, heterozygous female, and wild-type female mice (Fig. 4J–L). DPOAE measures at 8, 16, and 32 kHz were similarly comparable among groups (Fig. 4M–P), consistent with preserved cochlear function. A similar auditory phenotype was observed at 4 weeks of age (Supplementary Fig. 2). Heterozygous female mice also exhibited preserved rotarod performance compared with littermate controls at both 4 and 8 weeks of age (Fig. 4Q, Supplementary Fig. 2H). Together, these findings indicate that whole-body heterozygous *Kdm6a* deletion is insufficient to impair cochlear morphology or auditory function in young adult mice.

## 4. Discussion

Hearing impairment is commonly reported in KS2, but the underlying mechanisms remain poorly understood. Several clinical studies have described sensorineural hearing loss (SNHL), suggesting the possibility of intrinsic cochlear pathology [7–9]. However, recurrent otitis media, craniofacial abnormalities, and delayed middle-ear development are also common features of KS and may substantially contribute to conductive or mixed hearing loss [8]. This clinical heterogeneity makes it difficult to determine whether hearing impairment in KS2 primarily reflects cochlear dysfunction or secondary conductive abnormalities based on genotype–phenotype associations alone. To address this question, we generated and characterized two complementary *Kdm6a* deletion mouse models and performed genetic, physiological, and histological analyses to determine whether partial *Kdm6a* deficiency is sufficient to impair cochlear morphology and auditory function. Across the viable models examined, reduced *Kdm6a* dosage was not associated with overt cochlear pathology or baseline auditory dysfunction, suggesting that partial *Kdm6a* loss alone is insufficient to produce a robust primary cochlear phenotype in young mice.

In *Nestin-Cre; Kdm6a* conditional mutants, *Kdm6a* expression was reduced in bulk brain tissue but was not reduced in whole inner ear tissue; instead, whole-inner-ear *Kdm6a* mRNA levels were significantly higher than those in controls (Fig. 3A). Given the small sample size (n = 3 per group), the biological significance of this unexpected increase remains uncertain and requires confirmation in a larger cohort. *Nestin* expression in the inner ear is developmentally and spatially heterogeneous. During cochlear development, *Nestin* has been reported in the spiral ganglion region and in multiple cell populations of the organ of Corti, including developing hair and supporting cells, whereas its expression becomes more restricted in the mature cochlea [24]. More recent lineage-tracing studies have likewise identified only small populations of *Nestin*-expressing cells throughout the cochlear and vestibular tissues [25]. This heterogeneity may contribute to incomplete or nonuniform Cre-mediated recombination in cochlear cell populations, consistent with the persistent KDM6A immunoreactivity observed in our model. Because Cre-mediated recombination was not directly mapped in the spiral ganglion, auditory nerve, or central auditory nuclei, these findings should be interpreted as demonstrating reduced *Kdm6a* expression in bulk brain tissue without a corresponding reduction in inner ear *Kdm6a* mRNA expression or loss of cochlear KDM6A immunoreactivity, rather than as evidence that *Kdm6a* loss throughout the auditory pathway does not affect auditory function.

In the germline-derived global *Kdm6a* deletion model generated using *Sox2-Cre*, auditory function was similarly preserved in heterozygous female mice, a finding with particular clinical relevance because KS2 most commonly results from heterozygous loss-of-function variants in *KDM6A* [26, 27]. KDM6A immunoreactivity in cochlear tissues appeared lower in heterozygous females than in wild-type females; however, no overt cochlear pathology or baseline auditory dysfunction was observed. Thus, germline-derived heterozygous *Kdm6a* deletion was insufficient to produce an overt cochlear phenotype in young mice. These findings support the interpretation that partial *Kdm6a* loss alone may be insufficient to account for the hearing impairment observed in KS2 and highlight the potential contribution of conductive, mixed, developmental, central, or later-onset mechanisms

A notable difference between the *Nestin-Cre* and germline-derived heterozygous was that impaired rotarod performance was observed in *Nestin-Cre* conditional mutants but not in heterozygous females with global *Kdm6a* deletion. This difference may reflect differences in *Kdm6a* dosage and the spatial pattern of deletion between the two models. In the *Nestin-Cre* model, male *Nestin-Cre*; *Kdm6a^flox/Y^* mice lack a functional *Kdm6a* allele in Cre-expressing neural lineages, which may disrupt central nervous system development and contribute to impaired rotarod performance. In contrast, germline-derived heterozygous females retain one functional *Kdm6a* allele despite systemic deletion of the other allele. The absence of motor and auditory phenotypes in heterozygous females with global *Kdm6a* deletion suggests that partial *Kdm6a* loss is insufficient to disrupt early postnatal motor coordination or auditory function under the conditions examined. However, because mice with complete *Kdm6a* deletion were not recovered among viable offspring, we were unable to directly examine the effects of complete cochlear *Kdm6a* deficiency on auditory development and function. Therefore, we cannot exclude the possibility that complete loss of *Kdm6a* in cochlear tissues could impair cochlear development or auditory function.

Although *Kdm6a*/KDM6A was broadly expressed in the cochlea, the staining pattern was not uniform across cell types. KDM6A immunoreactivity was particularly prominent along the cell periphery of outer hair cells, whereas inner hair cells showed more evident nuclear and cytoplasmic signals. Supporting cells, auditory nerve fibers, spiral ganglion neurons, and the stria vascularis also showed detectable expression. These observations suggest that KDM6A expression in the inner ear is cell-type dependent. The functional significance of these cell-type-specific and subcellular staining patterns, particularly the peripheral enrichment observed in outer hair cells, remains unknown. Because KDM6A is primarily known as a chromatin-associated epigenetic regulator, the apparent non-nuclear localization is of particular interest but will require additional studies to determine its biological significance.

From a clinical perspective, our findings support a cautious reinterpretation of hearing impairment in KS2. Despite the reported occurrence of SNHL in individuals with KS2, partial *Kdm6a* loss in the viable mouse models examined did not produce overt cochlear pathology or baseline auditory dysfunction. These findings suggest that KDM6A haploinsufficiency alone may be insufficient to produce a robust primary cochlear phenotype and highlight the need to consider other mechanisms underlying KS2-associated hearing impairment. These may include conductive or mixed hearing loss associated with craniofacial and middle-ear abnormalities, as well as developmental, central, or later-onset auditory mechanisms. However, because mice with complete cochlear loss of *Kdm6a* were not recovered among viable offspring, our findings do not completely exclude a potential role for KDM6A in cochlear development or auditory function. Clinically, these results emphasize the importance of comprehensive audiologic and otologic evaluation in individuals with KS2 to distinguish cochlear from non-cochlear causes of hearing impairment and to identify potentially treatable middle-ear disease.

## 5. Limitations

This study has several limitations. First, we could not assess the effects of complete *Kdm6a* loss in postnatal inner ear tissues because mice with complete *Kdm6a* deletion were not recovered among viable offspring and because cochlear recombination was incomplete with the available Cre-driver lines. Second, species differences in inner ear development and susceptibility to middle-ear disease may limit the extrapolation of these findings to humans, particularly given the well-recognized contribution of otitis media to hearing outcomes in individuals with KS2. Third, we did not systematically evaluate middle-ear pathology, including infectious otitis media, in these mouse models. Although no overt abnormalities were identified by routine microscopic examination, additional studies are needed to determine whether conductive mechanisms contribute to hearing dysfunction in this setting. Fourth, quantitative cochlear *Kdm6a* mRNA analysis could not be performed in mice carrying the germline-derived heterozygous deletion because the relevant cochlear samples were no longer available. Therefore, the extent of *Kdm6a* reduction in the cochlea was not quantitatively confirmed. Finally, our analyses were limited to young mice evaluated at 4 and 8 weeks of age. We did not evaluate cochlear morphology, middle-ear pathology, or auditory function in middle-aged or older animals. Longitudinal studies will therefore be important to determine whether reduced *Kdm6a* dosage influences susceptibility to age-related hearing loss or other late-onset auditory phenotypes.

## 6. Conclusion

Across complementary mouse models, partial *Kdm6a* loss was insufficient to cause overt cochlear pathology or baseline auditory dysfunction in young mice. These findings provide *in vivo* functional evidence that reduced KDM6A dosage alone does not necessarily result in primary cochlear impairment and help refine the interpretation of hearing impairment associated with KS2. Rather than supporting a robust early-onset primary cochlear phenotype, our findings highlight the potential contribution of conductive, mixed, developmental, central, or later-onset mechanisms to KS2-associated hearing impairment.

Clinically, these findings highlight the importance of comprehensive otologic and audiologic evaluation in individuals with KS2, including careful assessment and management of middle-ear pathology and differentiation of cochlear from non-cochlear causes of hearing impairment. Future studies using alternative cochlear-specific genetic models or inducible postnatal deletion strategies will be important to directly determine whether complete cochlear loss of *Kdm6a* contributes to auditory dysfunction and to further define the role of KDM6A in inner ear development and maintenance.

## Data availability

Data are available from the corresponding author upon reasonable request.

## Supplementary Figures

**Supplementary Figure 1.**
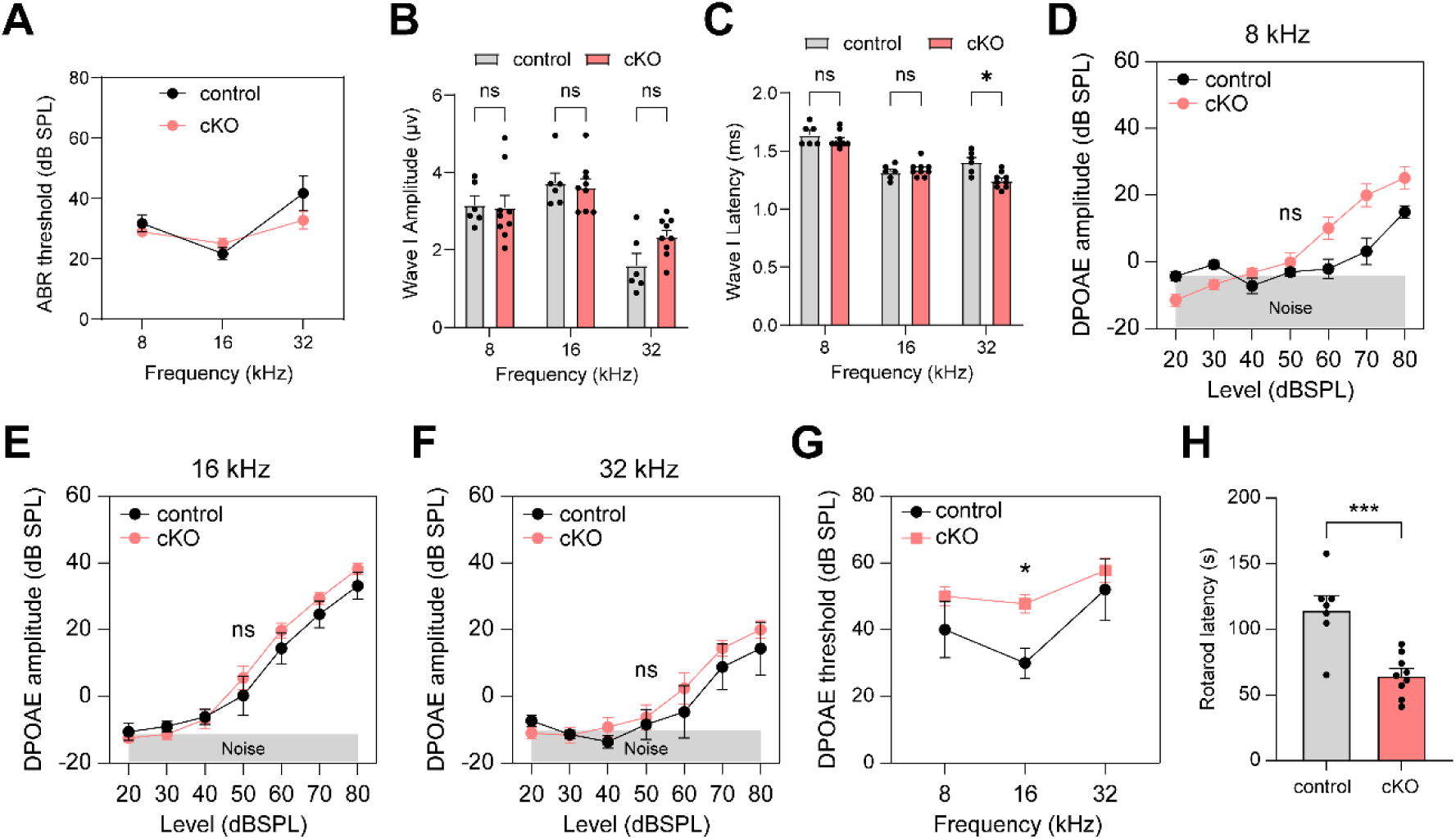
Assessment of Auditory Function and Rotarod Performance in *Nestin-Cre; Kdm6a* Conditional Mutant (cKO) Mice at 4 Weeks of Age. (A–C) ABR thresholds (A), wave I amplitudes (B), and wave I latencies (C) at 8, 16, and 32 kHz. (D–F) DPOAE amplitudes at 8, 16, and 32 kHz at 4 weeks. (G) DPOAE thresholds at 4 weeks of age. (H) Rotarod performance at 4 weeks of age. n = 5–7 in the control group and n = 9 in the cKO group; bars indicate mean ± SEM. *p < 0.05, ***p < 0.001; ns, not significant.

**Supplementary Figure 2.**
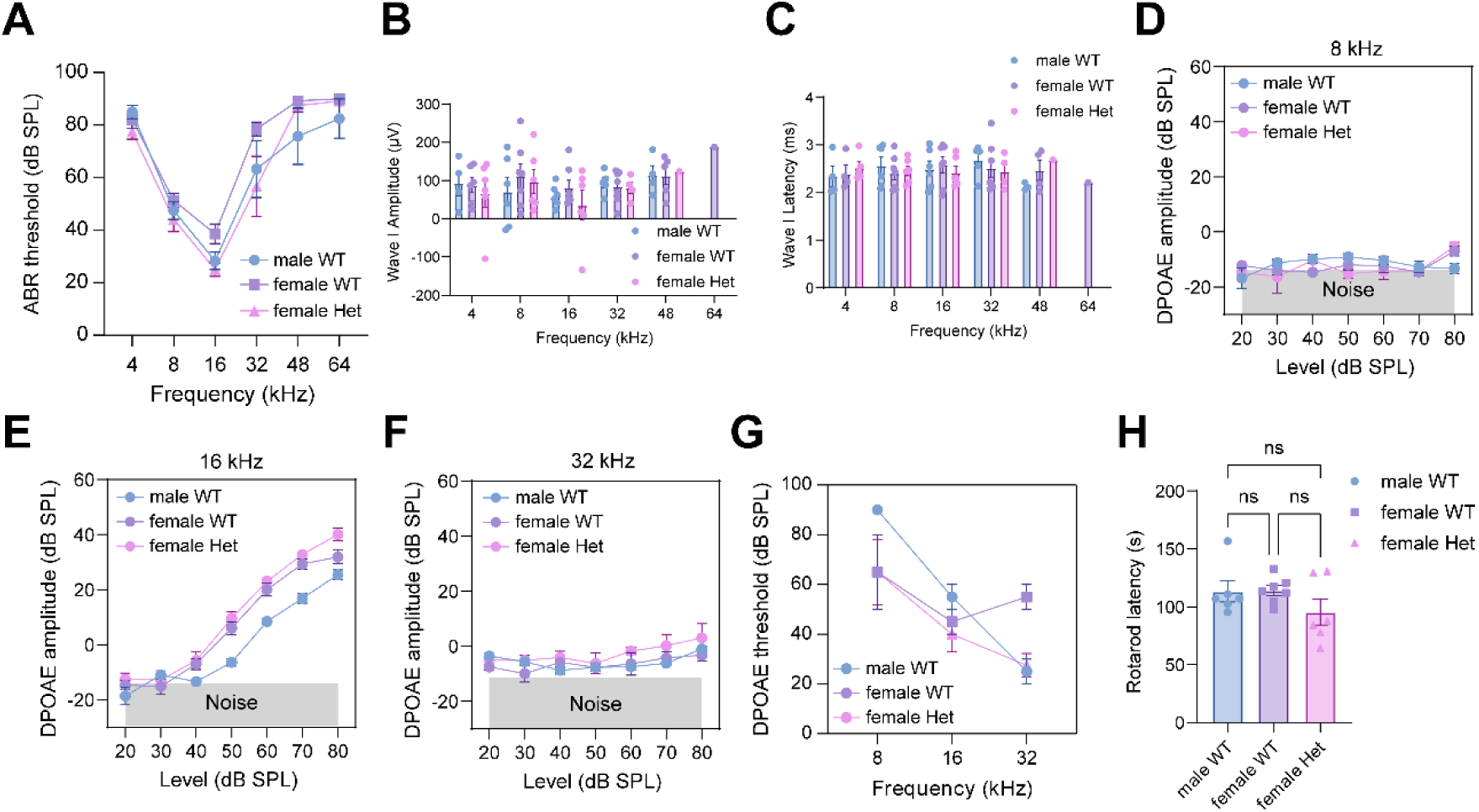
Assessment of Auditory Function and Rotarod Performance in Mice with Germline-Derived Heterozygous *Kdm6a* Deletion at 4 Weeks of Age. (A–C) ABR thresholds (A), wave I amplitudes (B), and wave I latencies (C) at 8, 16, and 32 kHz. (D–F) DPOAE amplitudes at 8, 16, and 32 kHz at 4 weeks of age. (G) DPOAE thresholds at 4 weeks of age. (H) Rotarod performance at 4 weeks of age. n = 6 for wild-type males, n = 6 for heterozygous females, and n = 7 for wild-type females. Bars indicate mean ± SEM. ns, not significant.

